# KIFC1 overexpression induces monopolar spindles by preventing centrosome separation during rapid cleavage divisions

**DOI:** 10.64898/2026.08.14.744973

**Authors:** Takahiro Yamamoto, Ai Kiyomitsu, Yang Ming, Tomomi Kiyomitsu

## Abstract

Bipolar spindle assembly is essential for accurate chromosome segregation. KIFC1, a conserved Ran-regulated minus-end-directed kinesin-14 motor, accumulates in the nucleus during interphase and promotes chromatin-mediated spindle assembly during mitosis and meiosis. In human oocytes, reduced KIFC1 levels destabilize meiotic spindles, a defect that can be rescued by increasing KIFC1 expression. However, how KIFC1 expression levels affect mitotic spindle stability during cleavage divisions in vertebrates remains unclear. Here, we show that whereas an approximately 50% reduction in KIFC1 causes no detectable defects in spindle assembly, approximately 10-fold overexpression of KIFC1 induces monopolar spindle formation, leading to chromosome mis-segregation and embryonic lethality in medaka early embryos. KIFC1 overexpression results in ectopic centrosomal localization during interphase, impairing the separation of duplicated centrosomes before mitotic entry. Analyses of KIFC1 mutants demonstrated that these centrosome separation defects require KIFC1’s microtubule-binding and motor activities and are further enhanced by deletion of KIFC1’s nuclear localization sequences. Together, our findings demonstrate that tight regulation of KIFC1 expression and its nuclear sequestration is essential for the proper separation and positioning of duplicated centrosomes before mitotic entry, thereby ensuring efficient bipolar spindle assembly during the rapid cleavage divisions of vertebrate embryos.

**Highlights:**

- KIFC1 accumulates in the nucleus and at the embryonic spindle midplane via the Ran pathway.
- Partial KIFC1 depletion does not impair spindle assembly in medaka early embryos.
- KIFC1 overexpression induces monopolar spindles by preventing centrosome separation.
- Centrosome separation defects require KIFC1 microtubule-binding and motor activity.

## Introduction

Bipolar spindle formation is essential for accurate chromosome segregation during meiosis and mitosis ^1,2^. Because the spindle is composed of numerous short, dynamic microtubules^3,4^, the nucleation and spatial organization of microtubules are fundamental to bipolar spindle assembly^2,5,6^. Previous studies have demonstrated that chromosomes generate a Ran-GTP gradient^7^, which activates spindle assembly factors (SAFs) including HURP^8,9^ and HSET/KIFC1^9–11^ by releasing inhibitory importins from SAFs in the vicinity of chromosomes. These SAFs stabilize microtubules near chromosomes^8,11,12^ and contribute to the formation of focused spindle poles^13^. In addition, recent studies have shown that Ran- GTP activates TPX2^14^ and augmin^15–17^ to promote microtubule-dependent branching microtubule nucleation around chromosomes. Together, these pathways coordinate bipolar spindle assembly, although their relative contributions and essentiality vary depending on the cellular context. For example, the Ran pathway is essential for acentrosomal spindle assembly during female meiosis^18–20^, likely because centrosomes are absent. In contrast, the Ran-GTP pathway is dispensable for spindle assembly in animal somatic cells^9^, probably because centrosome-mediated microtubule nucleation and the relatively small spindle size compensate for its loss^9^. We recently visualized spindle assembly processes during cleavage divisions in medaka (*Oryzias latipes*) early embryos (Fig. S1A) and demonstrated that embryonic spindle assembly requires the Ran-GTP pathway despite the presence of centrosomes to assemble specialized embryonic spindles^21^. However, how Ran-regulated spindle assembly factors cooperate with centrosome-mediated microtubule nucleation to assemble these specialized bipolar spindles during rapid cleavage divisions remains poorly understood.

KIFC1 (also known as HSET in mammals and XCTK2 in *Xenopus*) is a conserved Ran-regulated kinesin-14 motor that functions as non-processive minus-end-directed motor^22–24^. KIFC1 consists of three functional domains: an N-terminal tail domain, a coiled-coil stalk domain, and a C-terminal motor domain^24^. The N-terminal tail contains two microtubule-binding domains^25,26^, one of which contains a bipartite nuclear localization signal (NLSa and NLSb)^27^. The importin α/β complex recognizes these NLSs and transports KIFC1 into the nucleus while simultaneously inhibiting one of its microtubule-binding domains^26,27^. Importins are released from KIFC1 by Ran-GTP in the nucleus and near mitotic chromosomes^11,27^. The central stalk domain mediates KIFC1 homodimerization. *In vitro*, KIFC1 homodimers crosslink two microtubules through their N-terminal and C-terminal domains^28^ and generate inward sliding forces on antiparallel microtubules^29–31^. KIFC1 has also been implicated in the cross-linking and sliding of parallel microtubules^32–35^, which likely contributes to spindle pole focusing in diverse systems^10,13,36–42^. Notably, KIFC1 promotes clustering of supernumerary centrosomes in cancer cells^43^. In addition, a recent study reported that reduced KIFC1 levels destabilize meiotic spindles in human oocytes, a phenotype that can be rescued by restoring KIFC1 expression^44^. However, how KIFC1 expression levels influence centrosome separation and mitotic spindle stability in vertebrate embryos remains unclear.

In this study, we investigated localization and function of KIFC1 by modulating its protein level during cleavage divisions in medaka. We found that approximately 10-fold overexpression of KIFC1 causes abnormal spindle formation, leading to chromosome mis-segregation and embryonic lethality, whereas approximately 50% reduction in KIFC1 causes no detectable mitotic defects. KIFC1 overexpression induces ectopic centrosomal accumulation during interphase, preventing centrosome separation through its microtubule-binding and motor activities and thereby causing monopolar spindle formation. Our findings highlight the importance of proper centrosome separation and positioning before mitotic entry for efficient bipolar spindle assembly during rapid cleavage divisions of large vertebrate embryos.

## Results

### KIFC1 accumulates in the nucleus and around the metaphase spindle midplane via the Ran pathway in medaka early embryos

KIFC1 localization is regulated by Ran-GTP in cultured somatic cells^9,11,34^. To examine the intracellular distribution of KIFC1 during cleavage divisions in vertebrate embryos, we injected mRNA encoding mCherry-tagged KIFC1 (mCh-KIFC1) into one-cell-stage medaka (*Oryzias latipes*) embryos constitutively expressing EGFP-α-tubulin^21^ and analyzed its localization in four- and eight-cell-stage embryos. KIFC1 accumulates in the nucleus during interphase and at the microtubule-dense spindle midplane during metaphase (Fig. 1A, S1A). These localization patterns differed markedly from those of other mitotic kinesins. Eg5/KIF11, a homotetrameric kinesin-5 motor required for bipolar spindle assembly^45^, localized to centrosomes and the cytoplasm during interphase and was distributed throughout the mitotic spindle during metaphase (Fig. 1A). Kif2a, a kinesin-13 family microtubule depolymerase^46^, also localized to centrosomes and the cytoplasm during interphase, but became selectively enriched at spindle asters during metaphase (Fig. 1A), suggesting that microtubule motors functions at different locations during embryonic spindle assembly.

**Figure 1.**
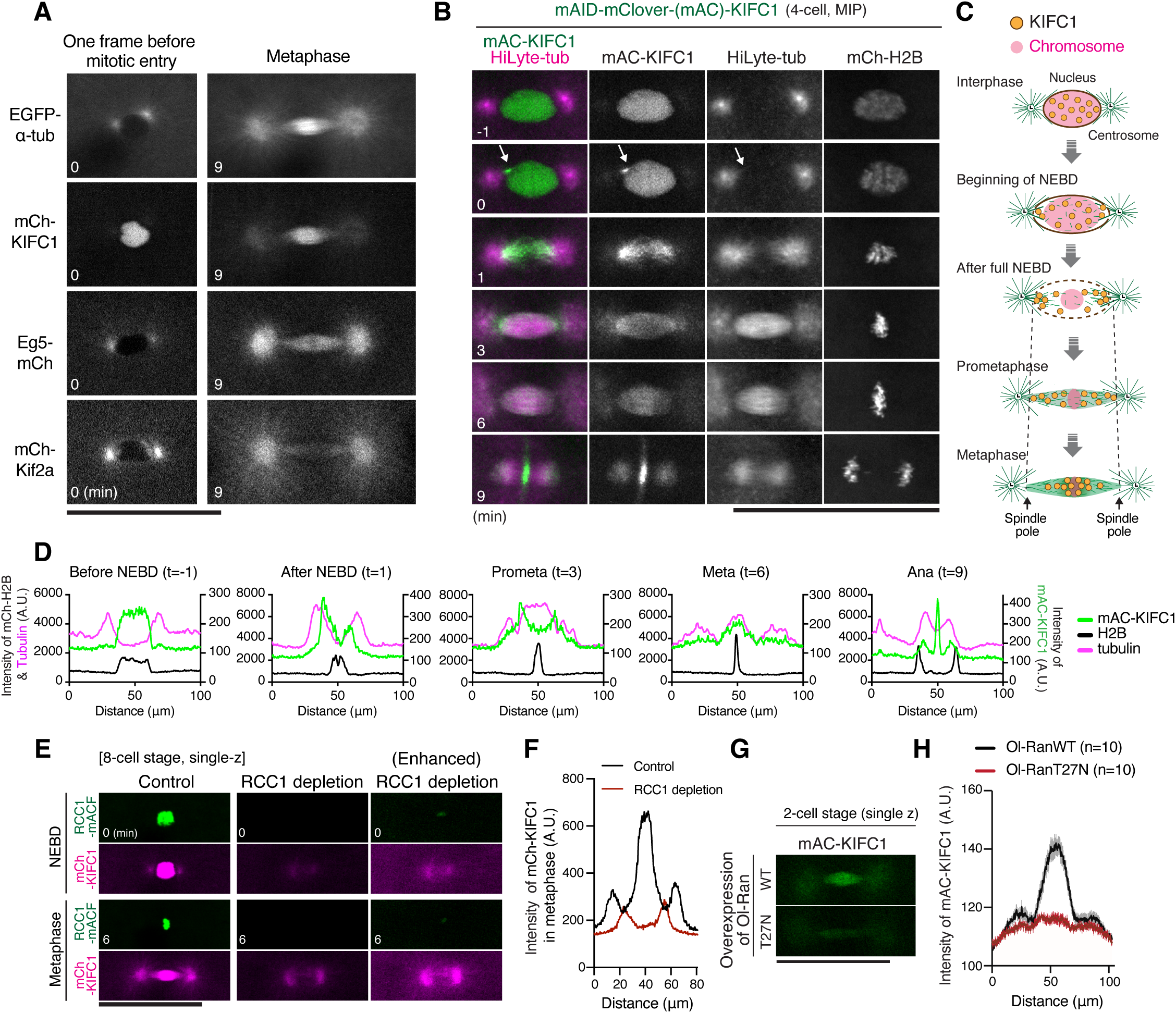
KIFC1 accumulates in the nucleus and at the spindle midplane via the Ran pathway during cleavage divisions. (A) Representative single-z section live images showing EGFP-α-tubulin and exogenously expressed mCherry-tagged motors one frame before mitotic entry (left) and at metaphase (right) in 4-cell-stage (tubulin and KIFC1) and 8-cell-stage (Eg5 and Kif2a) blastomeres. (B) Representative maximum-intensity-projection (MIP) live images showing mAID-mClover-KIFC1 (mAC-KIFC1), HiLyte 647-tubulin, and mCh-H2B in a 4-cell-stage blastomere. An arrow indicates a punctate KIFC1 signal. (C) Schematic representation of KIFC1 localization during spindle assembly in cleavage divisions. (D) Relative fluorescence intensity profiles from line scans of mAC-KIFC1 (green), mCh-H2B (black), and HiLyte 647- tubulin (magenta) across the spindles shown in (B). The left and right y-axes indicate the relative fluorescence intensities of H2B/tubulin and KIFC1, respectively. (E) Representative live images showing reduced nuclear localization of mCh-KIFC1 at NEBD (top) and reduced accumulation at the spindle midplane during metaphase (bottom) in RCC1-depleted 4-cell-stage blastomeres. (F) Line-scan analysis of mCh-KIFC1 fluorescence intensity across the metaphase spindle shown in (E). (G) Representative live images showing loss of endogenous mAC-KIFC1 accumulation at the metaphase spindle midplane in blastomeres expressing Ol-RanT27N (bottom), but not Ol-RanWT (control; top). (H) Line-scan analysis of mAC-KIFC1 fluorescence intensity across metaphase spindles in blastomeres expressing Ol-RanWT or Ol-RanT27N. Scale bars, 100 μm

To visualize endogenous KIFC1, we inserted an mAID-mClover (mAC) coding sequence at the N terminus of the KIFC1 gene using CRISPR/Cas9 (Fig. S1B)^21^. Homozygous knock-in medaka developed normally, and fertilized eggs obtained from homozygous pairs were used for live imaging. Endogenous mAC-KIFC1 also accumulated in the nucleus during interphase and around the metaphase spindle midplane in early embryos, including the one-cell stage (Fig. 1B, Fig. S1C). KIFC1 was evenly distributed throughout the nucleus before nuclear envelope breakdown (NEBD) (Fig. 1B). At the onset of NEBD, KIFC1 formed comet-like signals near the asters (Fig. 1B, Figs S1D and S1E) and then rapidly accumulated on both sides of the nuclear region adjacent to microtubule asters (Figs. 1B-1D), suggesting that nuclear KIFC1 rapidly associates with astral microtubules that invade the nucleus, forming a focused spindle pole-like structure. During prometaphase, KIFC1 signals gradually shifted from the spindle poles toward the spindle center (Fig. 1B, Fig. S1D) and eventually accumulated in the microtubule-dense region around the metaphase spindle midplane (Fig.1B-1D). After anaphase onset, KIFC1 remained weakly detectable on spindle microtubules (Fig. S1D) and also accumulated at spindle midzone (Fig. 1B). As cytokinesis progressed, KIFC1 localized to the cleavage furrow (Fig. S1F). KIFC1 exhibited similar nuclear and spindle localization in blastula stage embryos (Fig. S1G).

To test whether KIFC1 localization is regulated by the RCC1-Ran pathway, we next depleted RCC1 using auxin-inducible degron 2 (AID2) system^21^ . Exogenously expressed KIFC1 accumulated in the nucleus during interphase and around the spindle midplane during metaphase in control blastomeres but not in RCC1-depleted blastomeres (Figs. 1E and 1F). Consist with this result, expression of a dominant negative Ol-RanT27N mutant disrupted KIFC1 accumulation around the metaphase spindle midplane (Figs. 1G and 1H)^21^. These results indicate that KIFC1 undergoes dynamic changes in intracellular localization during spindle assembly and cytokinesis in medaka early embryos and that KIFC1 localization is regulated by Ran-GTP.

### Partial KIFC1 depletion has little effect on spindle assembly and embryogenesis in medaka

RCC1 depletion causes abnormal spindle formation followed by severe chromosome missegregation during cleavage in medaka^21^. To investigate the role of KIFC1 in embryonic spindle assembly, we next degraded endogenous KIFC1 using the AID2 system (Fig. 2A)^21^. We injected mRNAs encoding either mCherry-histone H2B (mCh-H2B) as a control or OsTIR1(F74G)-P2A-mCh-H2B to induce KIFC1 degradation into one-cell-stage embryos obtained from homozygous mAC-KIFC1 pairs. We subsequently performed live imaging in the presence of 5-Ph-IAA. Upon expression of OsTIR1(F74G)-P2A-mCh-H2B, mAC-KIFC1 fluorescence was reduced to approximately 50% of the control level by the four-cell stage (Figs. 2B and 2C). Normally, mAID-tagged proteins undergo further degradation as embryonic development proceeds owing to increasing OsTIR1 expression^21^. However, KIFC1 fluorescence remained at approximately 60% of the control level even at the eight- and sixteen-cell stages (Fig. S2A). Partial KIFC1 depletion did not cause obvious defects in bipolar spindle formation at the four-, eight-, or sixteen-cell stages (Figs. 2D, 2E, S2B), although it slightly increased the centrosome-to-centrosome distance at the four-cell stage (Figs. 2F). Consistent with these observations, partial KIFC1 depletion did not cause obvious defects in chromosome segregation (Fig. 2E) or embryogenesis (Fig. 2G). These results suggest that mitotic spindles assemble robustly and remain functional despite an approximately 50% reduction in KIFC1 during cleavage divisions in medaka early embryos.

**Figure 2.**
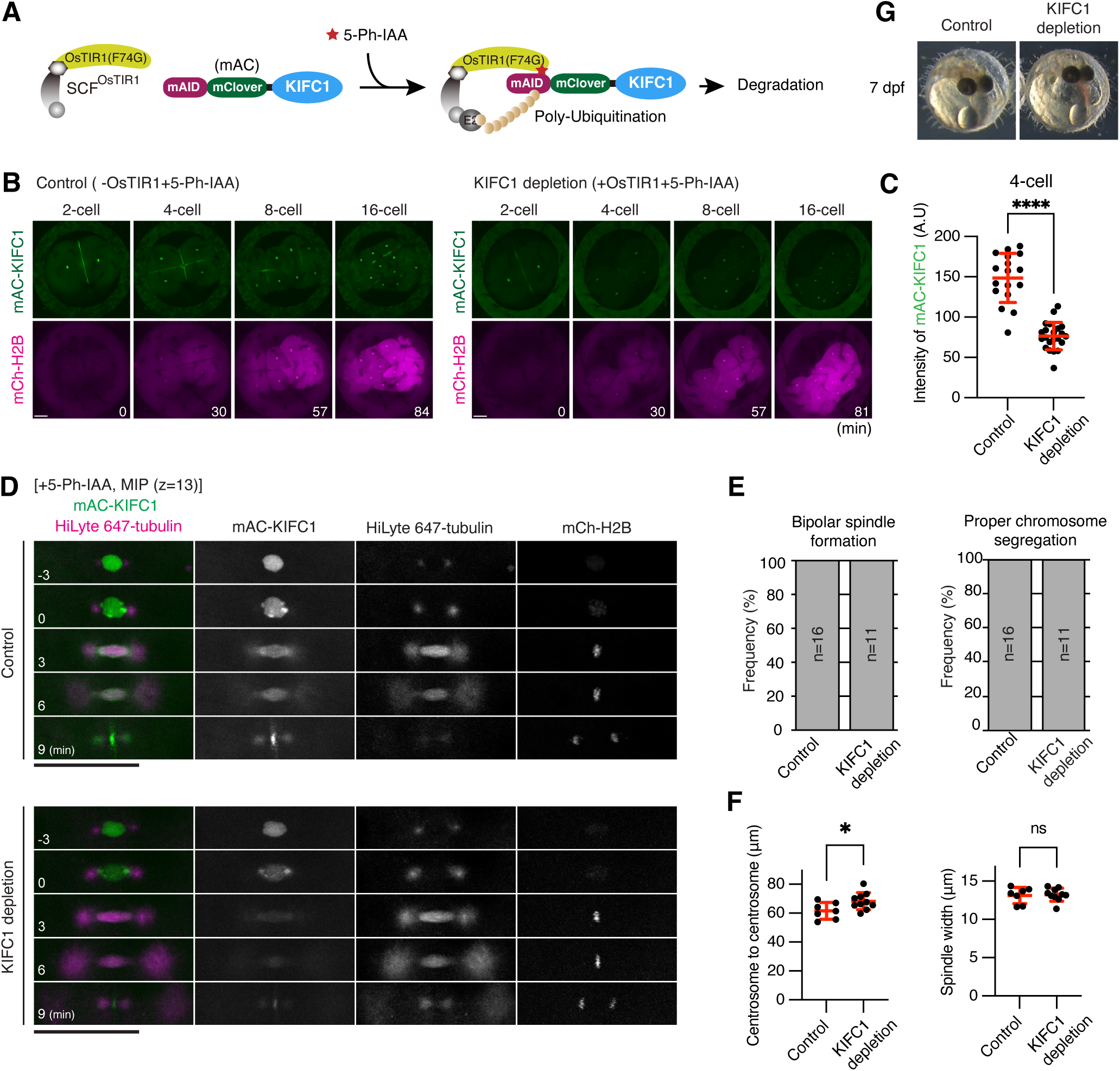
Partial KIFC1 depletion has little effect on spindle assembly and embryogeneis in medaka. (A) Schematic representation of auxin-inducible degron 2 (AID2)-mediated KIFC1 degradation. (B) Representative live images showing mAID-mClover-KIFC1 (mAC-KIFC1) and mCh-H2B fluorescence in control (left) and KIFC1-depleted embryos (right). OsTIR1(F74G) was not expressed in control embryos. (C) Quantification of mAC-KIFC1 fluorescence intensity in 4-cell-stage nuclei one frame before mitotic entry in control (n=17) and KIFC1-depleted blastomeres (n=22) from 5 and 7 embryos, respectively. Error bars indicate mean ± SD. Statistical significance was assessed using a two-sided Welch’s t-test. ****p < 0.0001. (D) Representative live images of control and KIFC1-depleted 4-cell-stage blastomeres showing mAC-KIFC1, mCh-H2B, and HiLyte 647-tubulin. The display ranges of mCh-H2B and HiLyte 647-tubulin were adjusted separately for control and KIFC1 depleted embryos because of differences in fluorescence intensity between the groups. (E) Frequencies of bipolar spindle formation (left) and proper chromosome segregation (right) during the 4-cell division. (F) Quantification of centrosome-to-centrosome distance (left) and spindle width (right) in control (n=7) and KIFC1-depleted (n=10) 4-cell-stage blastomeres from 5 and 4 embryos, respectively. Error bars indicate mean ± SD. Statistical significance was assessed using a two-sided Welch’s t-test. *p < 0.1. (G) Phase-contrast images of control and KIFC1-depleted embryos 7 days after mRNA injection. Scale bars, 100 μm.

### KIFC1 overexpression causes monopolar spindles and embryonic lethality in medaka embryos

We recently found that RCC1 overexpression induces spindle defects in a GEF activity-dependent manner in medaka early embryos (Yang Ming, unpublished results). To examine whether KIFC1 overexpression affects embryonic spindle assembly, we injected mRNAs encoding mCh-KIFC1 into EGFP-α-tubulin expressing embryos. Unexpectedly, KIFC1 overexpression induced bent and monopolar spindles in an expression-level dependent manner (Figs. 3A and B, Fig. S3A). These bent and monopolar spindles aligned the majority of chromosomes at metaphase (Fig. 3C). However, they subsequently caused abnormal chromosome segregation during anaphase (Fig. 3C). In addition, the paired asters failed to move apart, and cleavage furrows did not form between them (Figs. 3C and 3D), resulting in the formation of polyploid blastomeres. Despite these abnormalities, mitosis progressed without a delay (Fig. 3C, Fig. S3B), likely due to the absence of a functional spindle assembly checkpoint during medaka cleavage divisions^21^. These abnormal blastomeres underwent repeated abnormal divisions, resulting in embryonic lethality within 1 day post-fertilization (21/26 embryos) (Fig. 3E).

**Figure 3.**
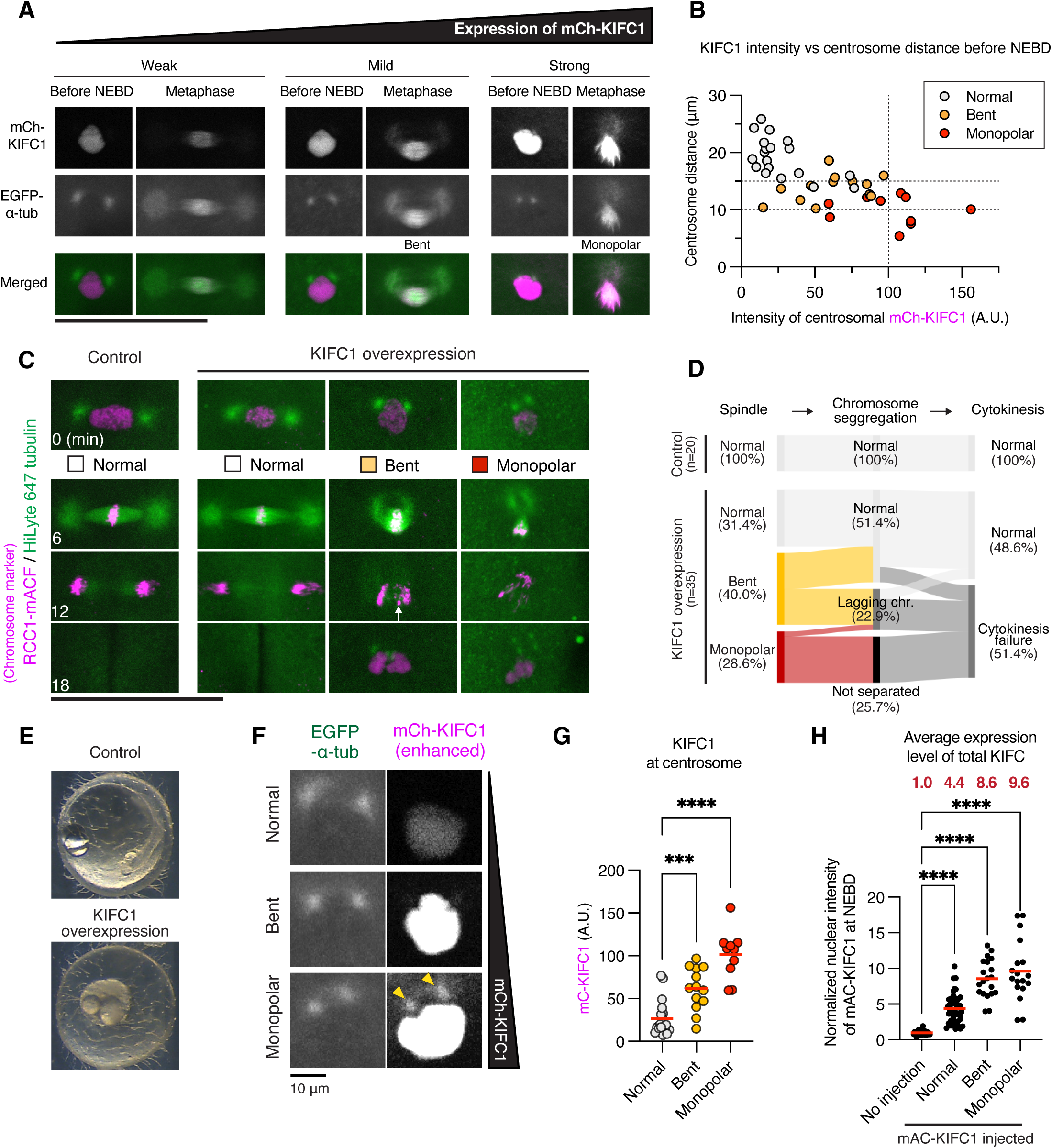
KIFC1 overexpression causes monopolar spindles and embryonic lethality. (A) Representative live images showing nuclei before NEBD and metaphase spindles in EGFP-α-tubulin-expressing blastomeres expressing different levels of mCh-KIFC1. The EGFP-α-tubulin display range was adjusted separately for KIFC1-overexpressing blastomeres because of differences in EGFP-α-tubulin expression levels among embryos. (B) Scatter plot of centrosome-to-centrosome distance and centrosomal mCh-KIFC1 intensity in blastomeres shown in Fig. S3C. (C) Representative live images showing chromosomes (RCC1-mAID-mClover-3xFLAG, RCC1-mACF) and spindles (HiLyte 647-tubulin) in control (left) and KIFC1-overexpressing (three columns on the right) blastomeres at the 4-cell stage. Bent and monopolar spindles cause chromosome segregation defects and cytokinesis failure. The white arrow indicates a lagging chromosome. The HiLyte 647-tubulin display range was adjusted separately. (D) Sankey diagrams showing the relationships among spindle formation defects, chromosome segregation defects, and cytokinesis failure during the 4-cell division. (E) Phase-contrast images of control (top) and mCh-KIFC1-expressing (bottom) embryos 1 day after mRNA injection. mCherry was overexpressed in control. (F) Representative live images showing EGFP-α-tubulin and mCh-KIFC1 one frame before NEBD. The yellow arrowheads indicate ectopic centrosomal localization of mCh-KIFC1. The EGFP-α-tubulin display range was adjusted separately. These images correspond to those at t = -6 in Fig. S3C. (G) Quantification of centrosomal mCh-KIFC1 fluorescence intensity in 4-cell-stage blastomeres. The mean KIFC1 fluorescence intensity at the two centrosomes was calculated for each blastomere. 19, 13, and 10 blastomeres were measured for normal, bent, monopolar spindles, respectively. Red bars indicate means. Statistical significance was assessed using One-way ANOVA with Dunnett’s multiple comparisons test. ***p < 0.001, ****p < 0.0001. (H) Quantification of normalized nuclear fluorescence intensity of total mAC-KIFC1 (endogenous plus ectopically expressed mAC-KIFC1) in blastomeres with different spindle morphologies. Red bars indicate means. Red numbers indicate the mean normalized fluorescence intensity for each group. Statistical significance was assessed using One-way ANOVA with Dunnett’s multiple comparisons test. ****p < 0.0001. Scale bars, 100 μm, except in (F), 10 μm.

To investigate the primary cause of the bent or monopolar spindles, we tracked centrosome behaviors before mitotic entry. During cleavage, centrosomes are normally duplicated and separated in cytoplasm after anaphase during the karyomere migration (Fig. S1A)^21,47,48^. Duplicated centrosomes subsequently associate with the nucleus and are positioned at opposite sides of the nucleus before NEBD (Fig. 3C)^21^. KIFC1 overexpression did not prevent centrosome-nucleus association but impaired centrosome separation and positioning before mitotic entry (Figs. 3B and 3C, Figs. S3C-E). Importantly, centrosome separation distance strongly correlated with spindle morphology: moderate separation (∼15 um) resulted in bent spindle, whereas limited separation (<10 um) produced monopolar spindles (Figs. 3B and 3C, Figs. S3C-E). We also found that KIFC1 ectopically accumulated at duplicated centrosomes during interphase in an expression-level dependent manner (Fig. 3F). Furthermore, increased centrosomal accumulation of KIFC1 correlated with defective centrosome separation and subsequent spindle assembly defect (Fig. 3G). To determine the threshold of KIFC1 overexpression required to induce spindle abnormalities, we injected mRNAs encoding mAC-KIFC1 into mAC-KIFC1 homozygous knock-in eggs (Fig. S3F) and quantified the total amount of KIFC1 (endogenous plus exogenous mAC-KIFC1) in cells exhibiting each phenotype. Cells expressing approximately 4.4-fold higher levels of nuclear KIFC1 were still be able to assemble bipolar spindles (Fig. 3H and Fig. S3F). In contrast, cells expressing approximately 8.6-fold or 9.6- fold higher levels of KIFC1, exhibited bent or monopolar-like spindles, respectively, although no clear threshold was evident (Fig. 3H). Together, these data suggest that KIFC1 overexpression allows KIFC1 to escape nuclear sequestration, leading to its ectopic accumulation at duplicated centrosomes, impaired centrosome separation during interphase, and ultimately abnormal spindle formation during early embryonic divisions.

### Ectopically localized KIFC1 prevents centrosome separation through its microtubule-binding and motor activities

To understand how ectopically localized KIFC1 at centrosomes impairs centrosome separation before mitotic entry, we generated five medaka (*Oryzias latipes*) KIFC1 mutants (Fig. 4A) and analyzed their localization and overexpression phenotypes. As expected, the NLS mutants, Ol-KIFC1 K10A R11A (NLSa mutant) and Ol-KIFC1 K29A K30A (NLSb mutant)^34^, partially or completely escaped nuclear sequestration and ectopically localized to centrosomes during interphase (Fig. 4B). Overexpression of these NLS mutants caused abnormal spindle formation (Fig. 4C), with higher frequencies than overexpression of Ol-KIFC1 WT (Fig. 4D, Fig. S3A). In contrast, Ol-KIFC1 Δ122, an N-terminal 122-amino-acid deletion mutant^49^, lacking both NLSs and the non-motor microtubule-binding site, did not cause spindle defects (Figs. 4C and 4D) although it also ectopically localized to centrosomes during interphase, similar to the NLS mutants (Fig. 4B). The C-terminal motor domain mutants, Ol- KIFC1 T380N (rigor-mutant)^49,50^ and N547K (motor-defective mutant)^34,49,51^, showed nuclear localization during interphase similar to wild-type KIFC1 (Fig. 4B), but neither mutant induced spindle defects (Figs. 4C and 4D), even when overexpressed to levels similar to those of wild-type KIFC1 (Fig. 4E).

**Figure 4.**
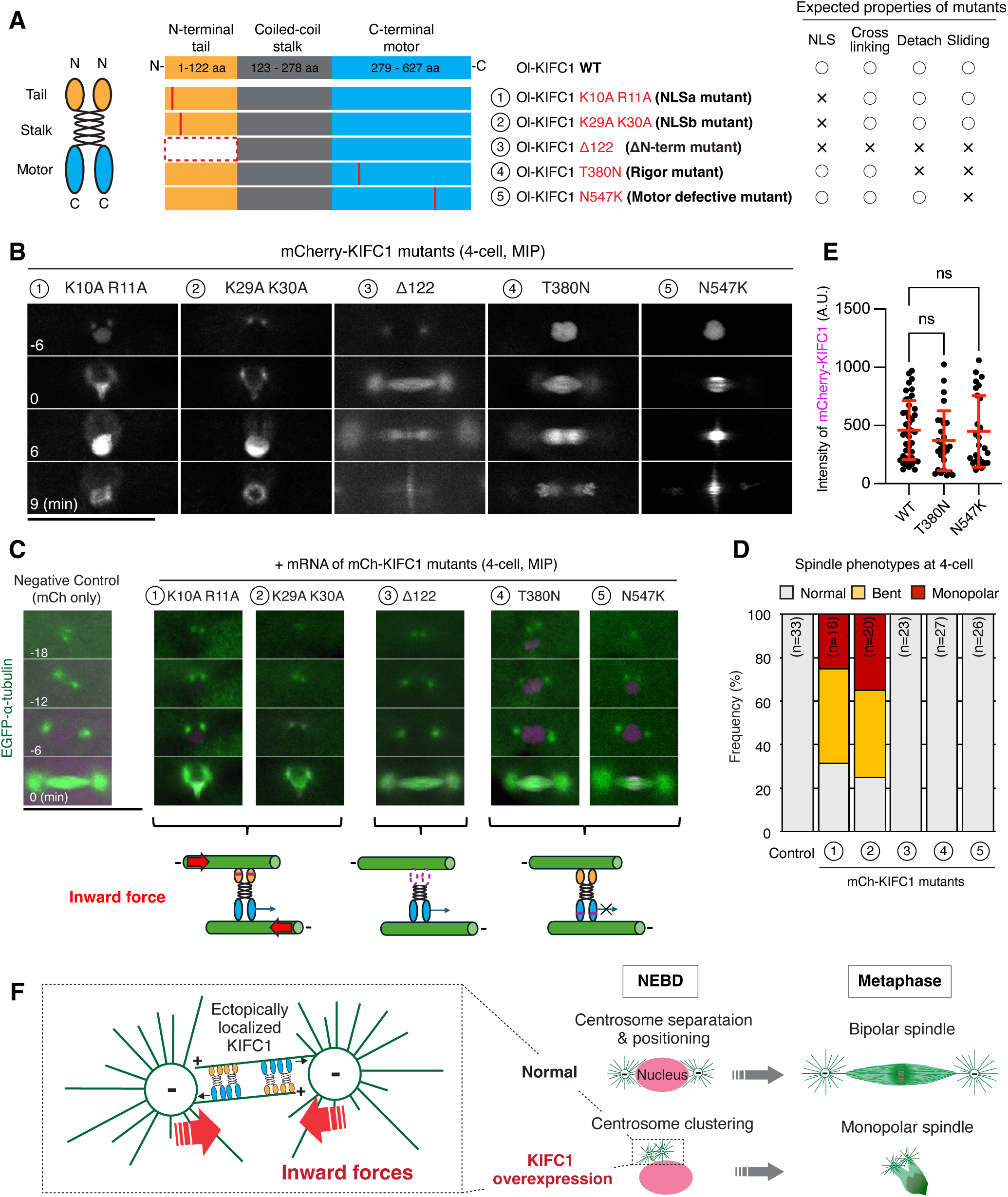
Ectopically localized KIFC1 prevents centrosome separation through its microtubule-binding and motor activities. (A) Schematic representation of the KIFC1 domain organization and five KIFC1 mutants with their expected properties. (B) Representative live images showing mCherry-tagged KIFC1 mutants during the 4-cell division. (C) Representative live images showing EGFP-α-tubulin in control and KIFC1 mutant-expressing embryos at the 4-cell stage. These images correspond to the same blastomeres shown in (B). EGFP-α-tubulin display range was adjusted separately. (D) Frequencies of spindle phenotypes during the 4-cell division following expression of the indicated KIFC1 mutants. (E) Quantification of nuclear mCh-KIFC1 fluorescence intensity for WT and two motor-domain mutants one frame before mitotic entry at the 4-cell stage. Error bars indicate mean ± SD. (F) Model illustrating how KIFC1 overexpression induces monopolar spindle formation by preventing centrosome separation before mitotic entry in medaka early embryo. See text for details. Scale bars, 100 μm.

Intriguingly, KIFC1 Δ122 (ΔN-terminal mutant) showed reduced accumulation around the spindle midplane at metaphase (Fig. 4B), suggesting that the N-terminal non-motor domain of KIFC1 is required for its accumulation around the spindle midplane. In addition, KIFC1 T380N (rigor mutant), which tightly and irreversibly binds to microtubules (Fig. 4A)^50^, failed to localize to the anaphase spindle midzone and cleavage furrow (Fig. 4B). Furthermore, its overexpression induced cytokinesis defects (Fig. S4A), likely by impairing the midzone localization of endogenous functional KIFC1 through heterodimerization (Fig. S4B).

Together, these results indicate that ectopic centrosomal localization of KIFC1 is not sufficient to impair centrosome separation but instead requires both its N-terminal non-motor microtubule- binding site and C-terminal motor activity (Fig. 4F). In addition, KIFC1 is required for cytokinesis and localizes to the anaphase spindle midzone through the dynamic microtubule-binding activity of its motor domain.

## Discussion

### Spatiotemporal localization and function of KIFC1 during cleavage divisions in medaka

In this study, we established a knock-in strain and investigated the spatiotemporal localization dynamics of endogenous KIFC1 and exogenously expressed KIFC1 mutants during cleavage divisions in medaka early embryos. We found that chromosome-derived Ran-GTP signals promote KIFC1 accumulation in the nucleus during interphase and at the specialized spindle midplane during metaphase in early embryos (Fig. 1), consistent with previous studies in other systems^11,34,41^. Our results indicate that Ran-mediated nuclear accumulation of KIFC1 serves at least two functions: promoting efficient bipolar spindle assembly during mitosis and sequestering KIFC1 away from centrosomes during interphase.

KIFC1 is transported into the nucleus by the importin-α/β complex, which recognizes the N-terminal NLSs of KIFC1 while inhibiting the overlapping microtubule-binding domain^11,26,27^. Upon entry into the nucleus, KIFC1 is released from importins by nuclear Ran-GTP. At the onset of NEBD, importin-free nuclear KIFC1 rapidly associates with astral microtubules that invade the nuclear region (Fig. 1B and Fig. S1E), likely together with other Ran-regulated SAFs such as HURP^8^ (Yang Ming, unpublished results). These SAFs subsequently crosslink and stabilize the astral microtubules to form a focused spindle pole-like structure (Fig. 1B). These microtubules likely provide a template for microtubule-dependent microtubule nucleation^3,15^, which is further promoted by Ran-dependent activation of augmin and TPX2 near chromosomes^16,17,52^. We propose a model in which sequential Ran-dependent activation of KIFC1 and other SAFs triggers a burst of oriented spindle microtubule nucleation from the spindle poles toward the chromosomes, thereby enabling rapid bipolar spindle assembly when centrosomes are positioned on opposite sides of the nucleus before mitotic entry (Figs. 1B, 1C, and 4F).

KIFC1 may function redundantly with other Ran-regulated SAFs and/or dynein during bipolar spindle assembly in early embryos^53^ (Yang Ming, unpublished results), which may explain why an approximately 50% reduction in KIFC1 has little or no detectable defects in spindle assembly. However, we cannot exclude the possibility that the remaining KIFC1 is sufficient to maintain spindle stability in early embryos. Unexpectedly, in contrast to the previous studies in *Drosophila*, Xenopus, and human cultured cells^10,41,54^, medaka KIFC1 accumulates at the spindle midzone during anaphase and cytokinesis in both early cleavage-stage and blastula-stage embryos (Fig. 1B, and Figs. S1D-G). This accumulation requires the dynamic microtubule-binding activity of the KIFC1 motor domain (Fig. 4B) and promotes cytokinesis in medaka embryos (Figs. S4A and 4B). Understanding how KIFC1 accumulates and functions at the spindle midzone during anaphase and cytokinesis and whether this role is specific to fish requires future investigation.

### KIFC1 overexpression causes monopolar spindles and embryonic lethality by preventing centrosome separation before mitotic entry

A key concept emerging from this study is that KIFC1 must be sequestered in the nucleus during interphase to ensure proper centrosome separation and positioning for bipolar spindle assembly during cleavage divisions. We found that approximately 10-fold overexpression allows KIFC1 to escape nuclear sequestration (Fig. 3H), resulting in its ectopic accumulation at centrosomes (Figs. 3F and 3G). This ectopic centrosomal KIFC1 subsequently prevents centrosome separation, likely by crosslinking and sliding antiparallel microtubules between duplicated centrosomes to generate inward pulling forces through its microtubule-binding and motor activities (Fig. 4F). A similar force-generation mechanism has been proposed for KIFC1/HSET-mediated clustering of supernumerary centrosomes in cancer cells^43^. Although centrosomal KIFC1 is unlikely to be activated by either nuclear Ran-GTP or the mitotic Ran-GTP gradient, it may nevertheless crosslink and slide antiparallel microtubules through the second microtubule-binding site within its N-terminal domain, which is not regulated by importins^26^.

In contrast to our findings in embryos, KIFC1 overexpression in HeLa cells causes elongated bipolar spindles but not monopolar spindles^34^, suggesting fundamental differences in both the mechanisms and mechanics of bipolar spindle assembly between small somatic cells and large, rapidly dividing embryonic cells. In canonical somatic cells, centrosomes are duplicated at the nuclear envelope during S-phase (Fig. S4C). The duplicated centrosomes remain physically linked until G2 phase but actively separate before mitotic entry through the prophase pathway, thereby facilitating bipolar spindle assembly^55^ (Fig. S4C). Interestingly, somatic cells have also evolved a prometaphase pathway, in which cells enter mitosis before complete centrosome separation and initially assemble monopolar-like spindles, which are subsequently converted into bipolar spindles following a short mitotic delay^56^. In sharp contrast, rapidly dividing fish embryos appear to rely exclusively on the prophase pathway. Centrosomes are duplicated immediately after mitosis and separate in the cytoplasm before associating with the nucleus, eventually becoming positioned on opposite sides of nucleus before NEBD^21,47^ (Fig. S4C). The prometaphase pathway is unlikely to operate during cleavage divisions in rapidly dividing vertebrate embryos because of the absence of a functional spindle assembly checkpoint (Fig. 3C)^21^. In addition, bursts of microtubule nucleation from two closely positioned centrosomes toward chromosomes preferentially generate bouquet-shaped monopolar spindles when the centrosome-to-centrosome distance is less than 10 μm (Figs. 3B and 3C, Figs. S3C and S3D). These bouquet-shaped monopolar spindles likely arise from the fusion of two initially independent microtubule arrays emanating from the duplicated centrosomes through extensive parallel microtubule bundling together with microtubule-dependent microtubule nucleation (Fig. S3D), thereby creating a physical linkage that prevents subsequent centrosome separation and the reorganization of monopolar spindles into bipolar spindles. Together, these observations suggest that centrosomes must be separated and positioned on opposite sides of the nucleus before mitotic entry to efficiently assemble bipolar spindles during the rapid cleavage divisions of large vertebrate embryos.

### Limitations of this study

Although we visualized the dynamic localization of endogenous KIFC1 in early embryos, it remains unclear how KIFC1 functions at the spindle pole-like region after NEBD, the metaphase spindle midplane, and the anaphase spindle midzone, and whether these spatiotemporal localization patterns and functions are conserved across vertebrates. In addition, our understanding remains limited regarding how KIFC1 expression is regulated and whether KIFC1 overexpression also induces monopolar spindles in blastula-stage or later embryos. Finally, several microtubule motors, including Eg5/kinesin-5, dynein, and KIFC3, have been reported to regulate centrosome separation during interphase in smaller somatic cells, but how these motors and other factors coordinate centrosome separation during the rapid cleavage divisions of large vertebrate embryos remains unclear and will require future investigation.

## Acknowledgments

We thank Toane Arata, Sumika Hagihara, Yoko Nakasone and the OIST animal resource section staffs for feeding and maintenance of medaka fish at OIST. We are grateful to NBRP Medaka (https://shigen.nig.ac.jp/medaka/) for providing OK-Cab (Strain ID: MT830).

## Author contributions

Conceptualization, TK; Investigation, TY, AK, and YM; Formal analysis, TY and YM; Methodology, AK and TK; Resources, AK and TK; Writing-original draft, TY; Writing-review & editing, TY, YM, and TK; Supervision, TK; Funding Acquisition, TK and AK.

## Data and materials availability

All data supporting findings of this study are available in the paper and its Supplementary Materials. All data of this study are stored at the corresponding author and available on reasonable request.

## Declaration of generative AI and AI-assisted technologies in the writing process

During the preparation of this work, the authors used ChatGPT (Open AI) to improve the language and readability of the manuscript. After using this tool, the authors reviewed and edited the content as needed and take full responsibility for the content of the published article.

## EXPERIMENTAL MODEL AND SUBJECT DETAILS

### Fish maintenance

Fish experiments were conducted in accordance with protocols (ACUP-2023-009, ACUP-2025-038, ACUP-2025-068) approved by the Animal Care and Use Committee at Okinawa Institute of Science and Technology Graduate University (OIST). The OK-Cab strain (MT830) of medaka (*Oryzias latipes*) was obtained from the National Bio-Resource Project Medaka (NBRP Medaka) and used as the parental strain. Medaka were raised and maintained as described previously^21^. Naturally fertilized eggs were collected from breeding pairs aged 2–9 months. Healthy fertilized eggs were used for imaging. Fluorescence of mAC-KIFC1 and RCC1-mACF was constantly observed in all knock-in embryos. In contrast, EGFP-α-tubulin fluorescence levels varied among embryos, as described previously^21^. Therefore, embryos with similar EGFP-α-tubulin fluorescence intensities were selected for all experiments.

## METHOD DETAILS

### Plasmid Construction

Donor plasmids and double-stranded DNAs (dsDNAs) for CRISPR/Cas9-mediated genome editing (Fig. S1B) were designed and constructed as described previously^21^. DNA fragments containing 5’ and 3’ homology arm (HA) regions were synthesized by Eurofins Genomics (Japan). mAID-mClover (mAC) cassette was inserted into the BamHI site. dsDNAs were amplified by PCR using modified primers containing a 5’ biotin modification and five consecutive 5’ phosphorothioate bonds (synthesized by Eurofins Genomics) and PrimeSTAR Max (Takara). Plasmids, medaka strains, and sequence information for the guide RNA (gRNA) and PCR primers used in this study are listed in Table S1, S2, S3, and S4, respectively.

### gRNA synthesis and in vitro transcription of mRNA

The gRNA was synthesized as described previously^21^. To synthesize Cas9 mRNA, the pCS2+hSpCas9 vector was linearized by NotI digestion, followed by in vitro transcription using the mMESSAGE mMACHINE SP6 Kit (Thermo Fisher Scientific, AM1340) according to the manufacturer’s instructions. The synthesized RNA was purified using an RNeasy Mini kit (Qiagen). To synthesize other mRNAs, template plasmids were linearized with NotI or BssHII.

### Microinjection

To exogenously express fluorescent fusion proteins, mRNAs at a concentration of 150 ng/µL were injected into one-cell-stage embryos as described previously^21^. Glass needles were prepared from borosilicate glass capillaries (Model No: G100F-4, Order No: 64-0787, Warner Instruments) using a needle puller (PC-100, Narishige), and attached to a capillary holder connected to a Femto Jet 4i microinjector (Eppendorf). The needles were manually controlled by a micro-manipulator (MN-153, Narishige) mounted on a stereomicroscope (Leica M80).

To simultaneously visualize microtubules together with green and red fluorescent proteins in early embryos, HiLyte 647-tubulin (Cytoskeleton, Inc. TL670M) was injected. HiLyte 647-tubulin (20 μg) were resuspended in 5 μL of RNase-free water. After centrifugation at 14,000 × g for 10 min at 4°C, the supernatant was collected, aliquoted, frozen in liquid nitrogen, and stored at -80°C. For co-injection of HiLyte 647-tubulin and mRNAs, a mixture containing final concentrations of 800 ng/μL HiLyte 647-tubulin and 150 ng/µL mRNA was injected using siliconized glass needles.

### Microscope system and live imaging

Live imaging was performed using a spinning-disc confocal microscope equipped with a 20× /0.95 NA water-immersion objective (APO LWD 20× WI λS; Nikon), 488-, 561-, and 640-nm lasers (Coherent), a CSU-W1 spinning-disk confocal unit (Yokogawa Electric Corporation), and an ORCA-Fusion sCMOS camera (Hamamatsu Photonics) mounted on an ECLIPSE Ti2-E inverted microscope with a Perfect Focus System (Nikon). Immersion water was automatically supplied to the objective using a water-immersion dispenser (Ti2-N-WID, Nikon). For imaging, embryos were mounted in a custom-made agarose chamber prepared in glass-bottom dishes (CELLview™, #627860, Greiner Bio-one). Seven #1.5 coverslips (18 mm×18 mm, 0.12–0.17 mm thickness; Matsunami) were stacked to generate a coverslip mold. Plastic molds with the same dimensions as the coverslip mold were fabricated using a 3D printer. These molds were used to form a shallow concave pocket in the agarose. After solidification, the coverslip mold was removed, producing a pocket approximately 0.8–1.2 mm wide and 3–5 mm deep on the glass surface. Four to nine embryos were aligned within the pocket with the blastodisc facing the glass bottom. The dishes were filled with approximately 2 mL of medaka balanced salt solution (BSS; 0.65% NaCl, 0.04% KCl, 0.02% MgSO_4_・7H_2_O, 0.02% CaCl_2_・2H_2_O; sterilized, pH 8.3, adjusted with 5% NaHCO_3_) ^18^, and imaging was performed at room temperature (24-25°C).

Z-stacks consisting of 13 optical sections at 5-μm intervals were acquired every 1 or 3 min with 1×1 camera binning. Green, red, and far red fluorescence images were acquired sequentially in this order, with exposure times of 500 ms, 1 s, 500 ms, respectively. X-Y-Z positions of 2–9 embryos were registered, and the embryos were automatically imaged for 6–10 h in a time-lapse experiment. For figures, 8-bit maximum-intensity-projection (MIP) images of z-stacks or single z-section images are shown as indicated. MIP images were generated using NIS-Elements. Signal intensities were linearly adjusted using Adobe Photoshop to optimize image clarity, and images were arranged using Adobe Illustrator. Phase-contrast images of embryos shown in Fig. 2G and Fig. 3E were acquired using a Leica M80 stereomicroscope equipped with a Leica MC190 HD camera.

### AID2-mediated protein degradation

AID2-mediated protein degradation was performed as described previously^21^. One-cell-stage embryos were injected with mRNAs encoding OsTIR1(F74G)-P2A-mCherry fusion proteins. After injection, embryos were gently agitated and cultured in BSS for 10–30 min. The medium was then replaced with 2 mL of BSS containing 10 μM 5-Ph-IAA. The embryos were transferred to the custom-made agarose chamber together with the treatment solution, oriented within the agarose pocket, and imaged using the spinning-disk confocal microscope described above.

### Quantification

For line-scan analyses of metaphase spindles, spindles with both centrosomes located within the same focal plane were selected (Fig. 1). To quantify the fluorescence intensities of EGFP-α-tubulin, mAC-KIFC1, and HiLyte 647-tubulin, line scan analyses were performed in Fiji (version 2.16.0/1.54p) or NIS-Elements (version 5.21.00, Nikon). A 5-pixel-wide line was drawn through both spindle poles and centrosomes (Fig. 1D). A 8-pixel-wide line was used for the analysis in Figs. 1F and 1H. Mean nuclear fluorescence intensities of mAC-KIFC1 (Figs. 2C, S2A and 3H) and mCh- KIFC1 (Fig. 4E) one frame before mitotic entry during the 4-cell division were measured using Fiji. Centrosome positions and mean fluorescence intensities of mCh-KIFC1 were measured using circular regions of interest (ROIs) with a diameter of 5.8 μm in Fiji (Figs. 3B, 3G, S3E). Centrosomal mCh-KIFC1 fluorescence intensities were measured two frames before initial spindle formation during the 4-cell division. EGFP-α-tubulin fluorescence was used to determine centrosome positions and define the ROIs. Fluorescence intensities in the neighboring cytoplasm were measured and subtracted as background in Figs. 2C, S2A 3B, 3G, S3E, 3H and 4E. Graphs were generated using Prism 10 (version 10.6.0, GraphPad Software, La Jolla, CA), SankeyMATIC, or Excel (version 16.105.1, Microsoft).

---

**Table S1:** Plasmids used in this study.

| No. | Name | Description | Reference | Related Figures |
| --- | --- | --- | --- | --- |
| 1 | pTK1077 | pCS2+mCherry2-OI-Kif2A | This study | Fig. 1A |
| 2 | pTK1126 | pCS2+OI-Eg5-mCherry2 | This study | Fig. 1A |
| 3 | pTK1042 | pCS2+mCherry2-OI-KIFC1 | This study | Fig. 1A, 1E-F, 3A-G, S3A-C, S3E, 4E |
| 4 | pTK1102 | OI-KIFC1-N: mAID-mClover | This study | Fig. S1B |
| 5 | pTK1064 | pCS2+OsTIR1(F74G)-P2A-mCherry2-OI-KIFC1 | This study | Fig. 1E-F |
| 6 | pTK1035 | pCS2+mCherry2-OI-RanT27N | Kiyomitsu et al., 2024 | Fig. 1G-H |
| 7 | pTK1040 | pCS2+mCherry2-OI-RanWT | Kiyomitsu et al., 2024 | Fig. 1G-H |
| 8 | pTK1076 | pCS2+_mCherry2-H2B | Kiyomitsu et al., 2024 | Fig. 1B, 1D, S1D, 2B-G, S2A-B |
| 9 | pTK1073 | pCS2+mCherry2-α-tubulin | Kiyomitsu et al., 2024 | Fig. S1G |
| 10 | pTK1063 | pCS2+miRFP670nano3-H2B | Kiyomitsu et al., 2024 | Fig. S1G |
| 11 | pTK1050 | pCS2+OsTIR1(F74G)-P2A-mCh-H2B | Kiyomitsu et al., 2024 | Fig. 2B-G, S2A-B |
| 12 | pTK1075 | pCS2+mCherry2 | This study | Fig. 3C-E, S3A-C, S3E, 4B |
| 13 | pTY19 | pCS2+mcherry2-OI-KIFC1_delta N term | This study | Fig. 4B-D |
| 14 | pTY20 | pCS2+mcherry2-OI-KIFC1 T380N rigor mutant | This study | Fig. 4B-E, S4A-B |
| 15 | pTY21 | pCS2+mcherry2-OI-KIFC1_T547K_motor_defective_mutant | This study | Fig. 4B-E |
| 16 | pTY25 | pCS2+mCherry2-OI-KIFC1 K10A R11A NLSa mutant | This study | Fig. 4B-D |
| 17 | pTY26 | pCS2+mCherry2-OI-KIFC1 K29A K30A NLSb mutant | This study | Fig. 4B-D |
| 18 | pTY28 | pCS2+mAID-mClover-OI-KIFC1 | This study | Fig. 3H, S3F |

**Table S2:** Medaka strains used in this study.

| No. | Name | Description | Plasmids used or references |
| --- | --- | --- | --- |
| 1 | OK-Cab (MT830) | National Bio-Resource Project Medaka | Kiyomitsu et al., 2024 |
| 2 | EGFP-α-tubulin | A knock-in strain having EGFP-α-tubulin at the tubulinα-1B locus | Kiyomitsu et al., 2024 |
| 3 | RCC1-mACF | A knock-in strain having mAID-mClover-3xFLAG (mACF) at the RCC1 locus. | Kiyomitsu et al., 2024 |
| 4 | mAC-KIFC1 | A knock-in strain having mAID-mClover (mAC) at the KIFC1 locus. | pTK1102, pCS2+hSpCas9 |

**Table S3:**
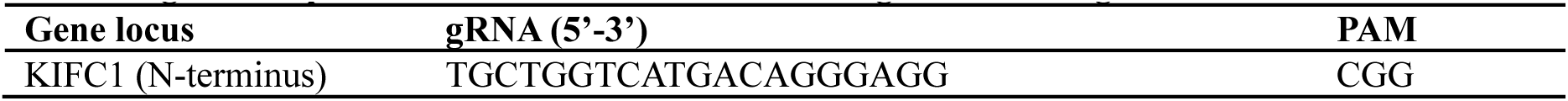
gRNA sequences for CRISPR/Cas9-mediated genome editing.

**Table S4:**
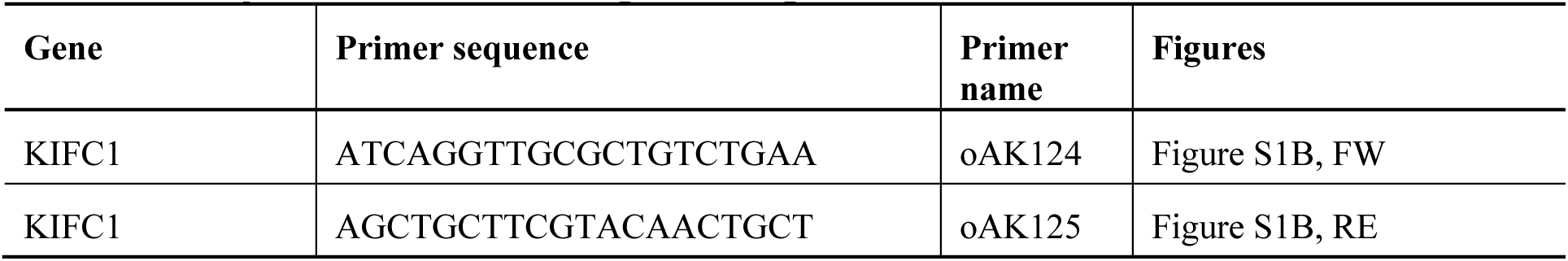
PCR primers used to confirm gene editing.

**Figure S1.**
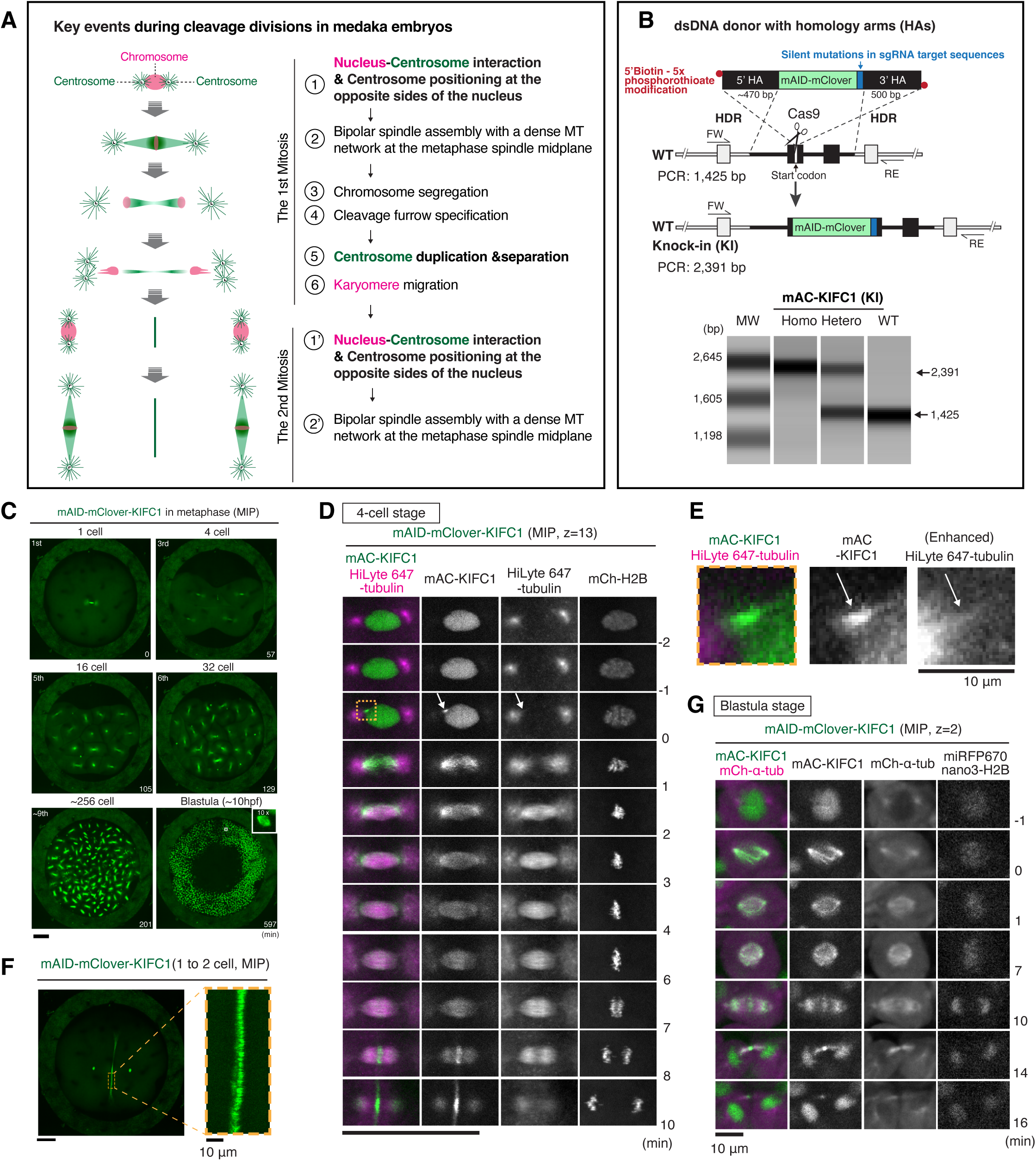
Establishment of mAID-mClover-KIFC1 knock-in strain and live imaging of endogenous KIFC1 during eary embryonic divisions, related to Figure 1. (A) Diagram showing key events during cleavage divisions in medaka embryos. (B) Top, schematic representation of the strategy used to generate the mAC-KIFC1 knock-in (KI) strain using dsDNA as a donor. Bottom, PCR-based genotyping of the KIFC1 locus in the parental wild-type (WT) and KI strains. A single band of approximately 2.4 kb confirms homozygous insertion in the KI strain. (C) Representative maximum-intensity-projection (MIP) live images showing metaphase spindles at the indicated embryonic stages. (D) Representative live images showing mAC-KIFC1, HiLyte 647-tubulin, and mCh-H2B in a 4-cell-stage blastomere. The arrow indicates a punctate KIFC1 signal. The boxed region is enlarged in (E). (E) Enlarged live images showing colocalization of mAC-KIFC1 and HiLyte 647-tubulin in the boxed region in (D). (F) Representative live image showing KIFC1 accumulation at the cleavage furrow. Right, enlarged image showing KIFC1 localization along putative antiparallel microtubules at the cleavage furrow. (G) Representative live images showing mAC-KIFC1, mCh-α-tubulin and miRFPnano3-H2B in a blastula-stage blastomere. Scale bars, 100 μm, except in (E-G), 10 μm.

**Figure S2.**
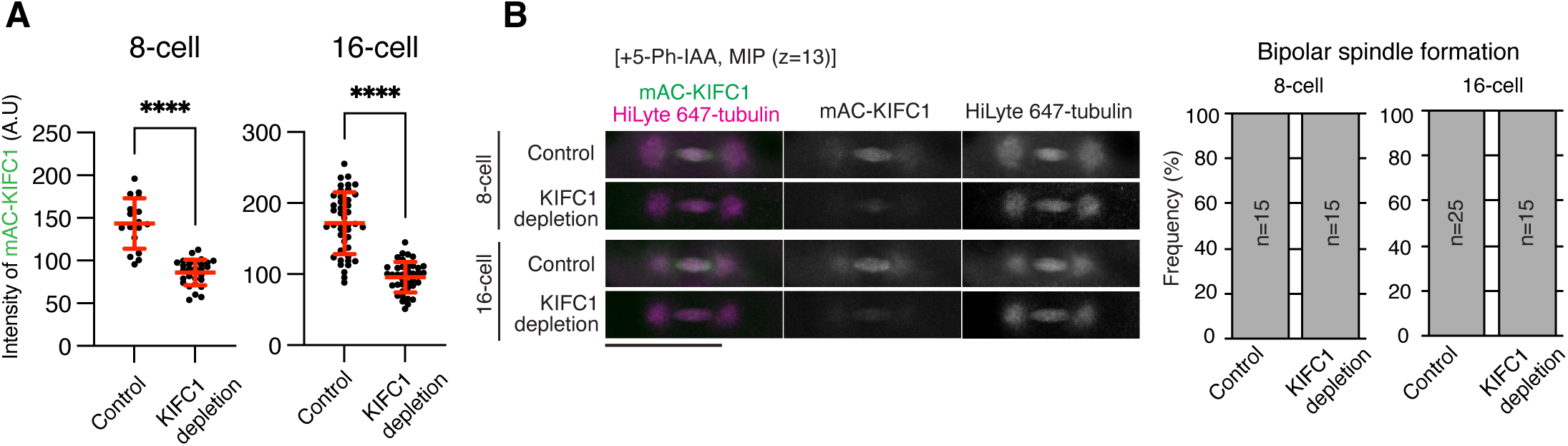
Partial KIFC1 depletion has little effect on bipolar spindle formation in 8-cell- and 16-cell-stage blastomeres, related to Figure 2. (A) Quantification of mAC-KIFC1 fluorescence intensity in nuclei at the 8-cell and 16-cell stages one frame before mitotic entry in control (8-cell: n = 17 from 4 embryos; 16-cell: n = 43 from 5 embryos) and KIFC1-depleted blastomeres (8-cell: n = 29 from 7 embryos; 16-cell: n = 39 from 6 embryos). Error bars indicate mean ± SD. Statistical significance was assessed using two-sided Welch’s t-tests. ****p < 0.0001. (B) Left, representative live images of control and KIFC1-depleted blastomeres at the 8-cell and 16-cell-stages showing mAC-KIFC1, mCh-H2B, and HiLyte 647-tubulin. The HiLyte 647-tubulin display range was adjusted separately. Right, frequencies of bipolar spindle formation in control and KIFC1-depleted blastomeres at the 8-cell and 16-cell stages. Scale bars, 100 μm.

**Figure S3.**
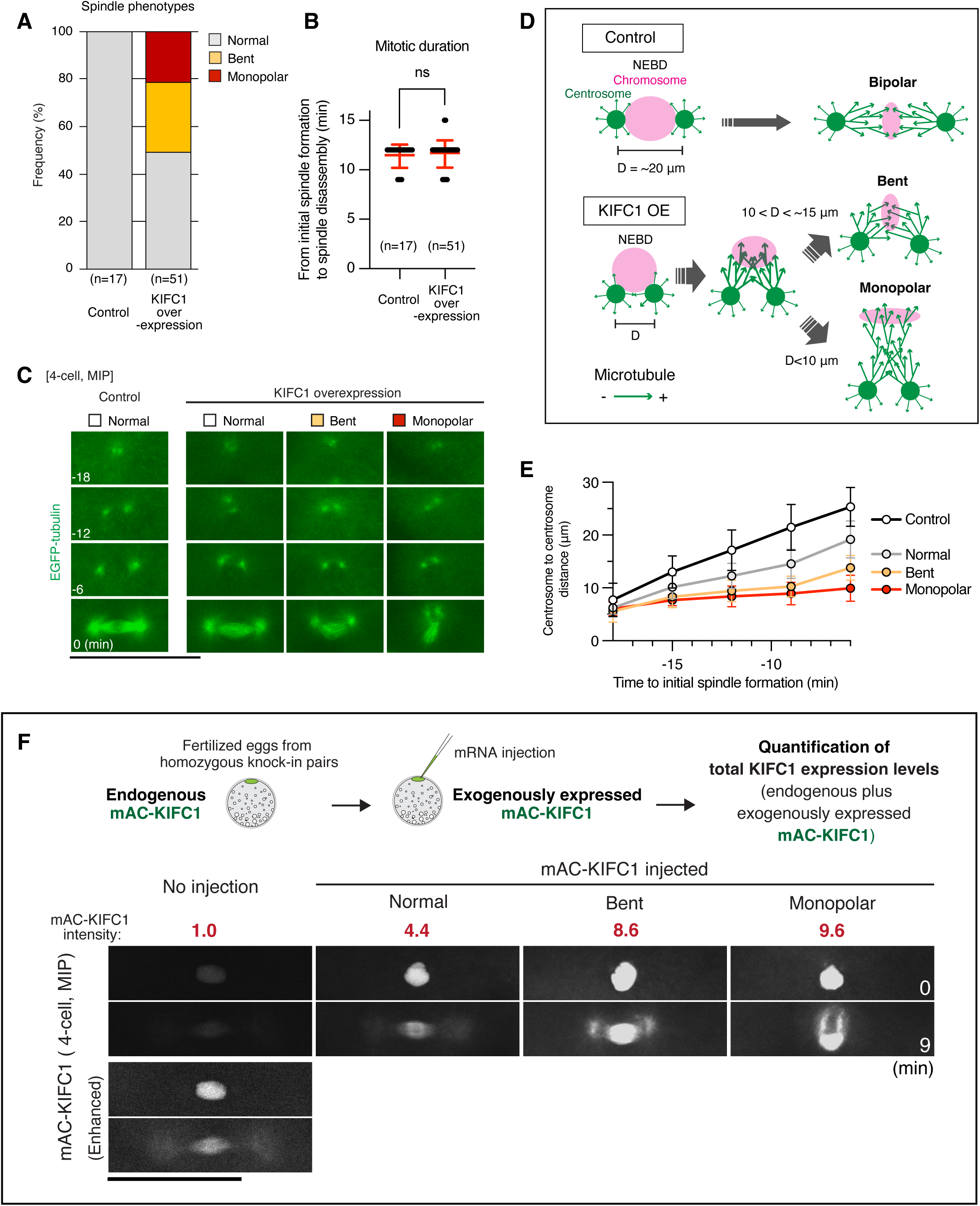
Approximately 10-fold overexpression of KIFC1 causes monopolar spindle formation by preventing centrosome separation, related to Figure. 3. (A) Frequencies of spindle phenotypes during the 4-cell division. (B) Scatterplots of mitotic duration in control (11.5 ± 1.2 min, n = 17) and mCh-KIFC1 expressing (11.7 ± 1.4 min, n = 51) blastomeres at the 4-cell stage. Mitotic duration was measured from initial spindle formation to spindle disassembly. (C) Representative live images of EGFP-α-tubulin showing centrosome separation in control (left) and mCh-KIFC1-expressing (three columns on the right) 4-cell-stage blastomeres. The EGFP-α-tubulin display range was adjusted separately for KIFC1-overexpressing blastomeres to improve visualization of spindle morphology because of differences in EGFP-α-tubulin expression levels among embryos. (D) Diagrams showing the relationship between centrosome separation with spindle morphology following KIFC1 overexpression. (E) Graphs showing changes of centrosome-to-centrosome distance in control blastomeres (black, n =14) and mCh-KIFC1-expressing blastomeres that formed bipolar (gray, n = 19), bent (yellow, n = 13), or monopolar (red, n = 10) spindles. (F) Diagrams illustrating the method used to quantify total KIFC1 expression levels following exogenous expression of mAC-KIFC1 (top). Representative live images showing nuclei before NEBD and metaphase spindles in blastomeres expressing different levels of mAC-KIFC1 (bottom). Red numbers indicate the mean normalized mAC-KIFC1 fluorescence intensity for each group. Scale bars, 100 μm.

**Figure S4.**
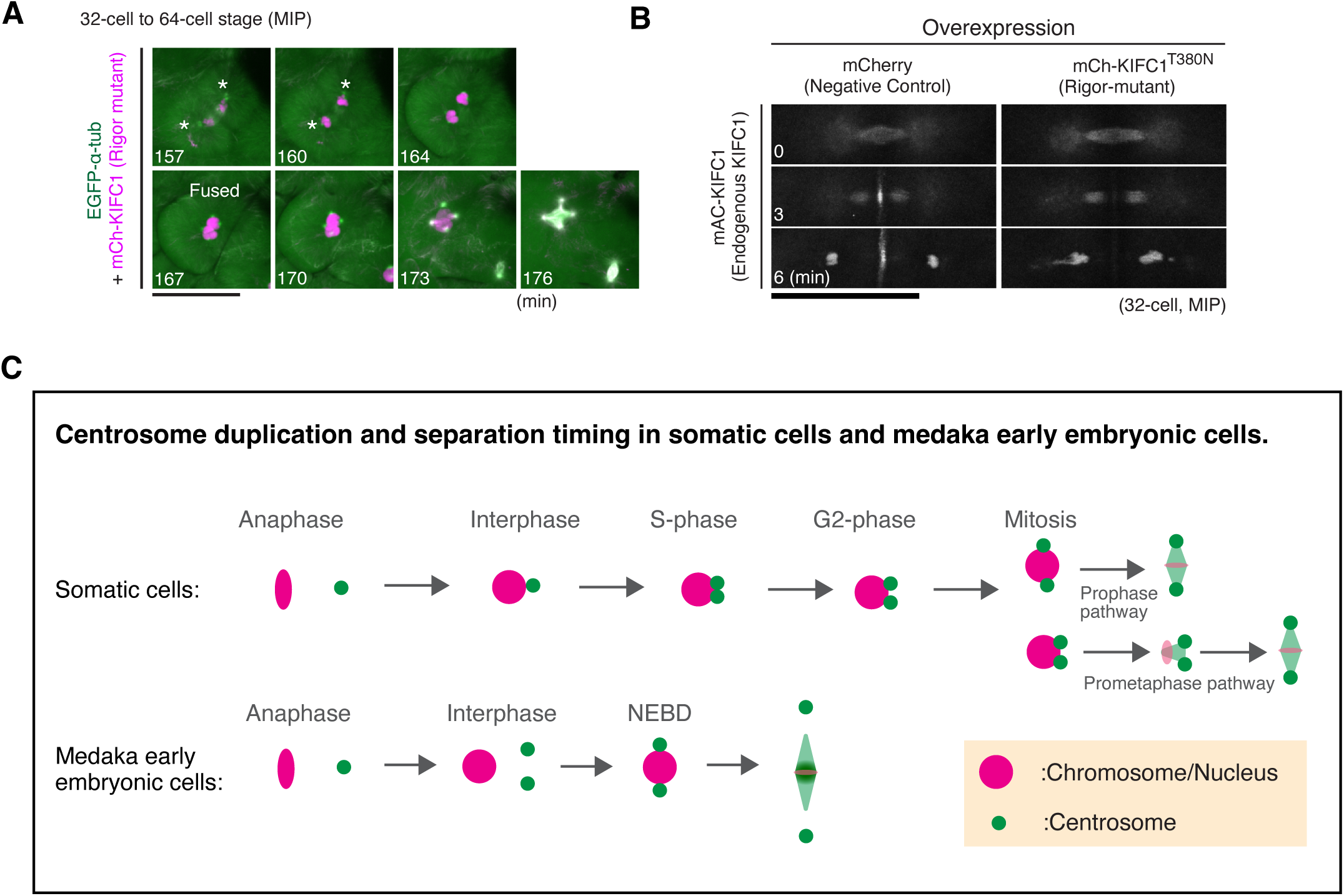
Expression of the KIFC1 T380N rigor mutant induces cytokinesis defects, related to Figure 4. (A) Representative live images showing EGFP-α-tubulin and mCh-KIFC1 T380N from 32-cell to 64-cell stage. Following cytokinesis failure, two separated daughter nuclei (asterisks) subsequently fused and formed an abnormal multipolar spindle at the next mitosis. (B) Representative live images showing endogenous mAC-KIFC1 in control (mCherry expressing) and mCh-KIFC1 T380N-expressing blastomeres at the 32-cell stage. Expression of mCh-KIFC1 T380N reduces endogenous KIFC1 localization to the cleavage furrow. (C) Diagrams comparing the timing of centrosome duplication and separation in somatic cells and medaka early embryonic cells. In somatic cells, centrosomes duplicate and separate at the nuclear envelope, whereas in medaka early embryos, they duplicate and separate in the cytoplasm. In addition, somatic cells use both prophase and prometaphase pathways for centrosome separation, whereas medaka early embryonic cells appear to rely primarily on the prophase pathway, with no detectable prometaphase pathway. Scale bars, 100 μm. separation before mitotic entry in medaka early embryo. See text for details. Scale bars, 100 μm.

